# Large frugivores matter more on an island: insights from island-mainland comparison of plant-frugivore communities

**DOI:** 10.1101/2020.07.31.229278

**Authors:** Rohit Naniwadekar, Abhishek Gopal, Navendu Page, Sartaj Ghuman, Vivek Ramachandran, Jahnavi Joshi

**Affiliations:** Nature Conservation Foundation, 1311, “Amritha”, 12^th^ Main, Vijayanagar 1^st^ Stage, Mysuru, Karnataka, India 570017; Wildlife Institute of India, Post Box no. 18, Chandrabani, Dehradun, Uttarakhand, India 248001; National Centre for Biological Sciences, Tata Institute of Fundamental Research, Bellary Road, Bengaluru, Karnataka, India 560065; Centre for Cellular and Molecular Biology, Uppal Road, Hyderabad, Telangana, India 500007

**Keywords:** community phylogenetics, insular communities, Narcondam Hornbill, plant-frugivore interactions, seed dispersal

## Abstract

Endozoochory, a mutualistic interaction between plants and frugivores, is one of the key processes responsible for maintenance of tropical biodiversity. Islands, which have a smaller subset of plants and frugivores when compared with mainland communities, offer an interesting setting to understand the organization of plant-frugivore communities vis-a-vis the mainland sites. We examined the relative influence of functional traits and phylogenetic relationships on the plant-seed disperser interactions on an island and a mainland site. The island site allowed us to investigate the organization of the plant-seed disperser community in the natural absence of key frugivore groups (bulbuls and barbets) of Asian tropics. The endemic Narcondam Hornbill, was the most abundant frugivore on the island and played a central role in the community. Species strength, a measure of relevance of frugivores for plants, of frugivores was positively associated with their abundance. Among plants, figs had the highest species strength and played a central role in the community. Island-mainland comparison revealed that the island plant-seed disperser community was more asymmetric, connected and nested as compared to the mainland community. Neither phylogenetic relationships or functional traits (after controlling for phylogenetic relationships) were able to explain the patterns of interactions between plants and frugivores on the island or the mainland pointing towards the diffused nature of plant-frugivore interactions. The diffused nature is a likely consequence of plasticity in foraging behavior and trait convergence that contribute to governing the interactions between plants and frugivores.

## INTRODUCTION

Endozoochory, a mutualistic interaction between plants and frugivorous animals, represents a critical stage in plant regeneration and contributes to the maintenance of tree diversity in tropics (Terborgh et al. 2002). The community of fleshy-fruited plants and seed dispersers form a network of bipartite interactions. These networks could be influenced by different ecological (e.g. functional traits, phenological overlap) and evolutionary (e.g. co-evolution) processes (Vázquez et al. 2009; Guimarães Jr et al. 2011) whose relative roles are yet to be sufficiently investigated.

Shared evolutionary history can play a role in governing the interactions between plants and frugivores. Traits relevant to the seed dispersal such as beak width have been demonstrated to exhibit a phylogenetic signal (Rezende et al. 2007). On the other hand, trait convergence could result in unrelated species interacting with similar partners (Peralta 2016). Thus, interactions could be influenced by trait complementarity – which itself could be modulated by trait conservatism – or by trait convergence (Rezende et al. 2007; Bastazini et al. 2017).

To assess the relative roles of functional traits and their phylogenetic history in determining patterns of ecological interactions, Bazatazini et al. (<u>2017)</u> present an ecophylogenetic approach which outlines four scenarios based on the relationship between the degree of phylogenetic conservatism in species traits and their phylogenetic history. The four scenarios enable determining whether the interactions between plants and frugivores are 1) mediated by traits (with no trait conservatism), 2) mediated by phylogeny (with trait conservatism), 3) mediated by phylogeny (with no trait conservatism), or 4) mediated by both traits and phylogeny (with some trait conservatism) (see Fig. 1 in Bastazini et al. 2017 for additional details).

**Figure 1.**
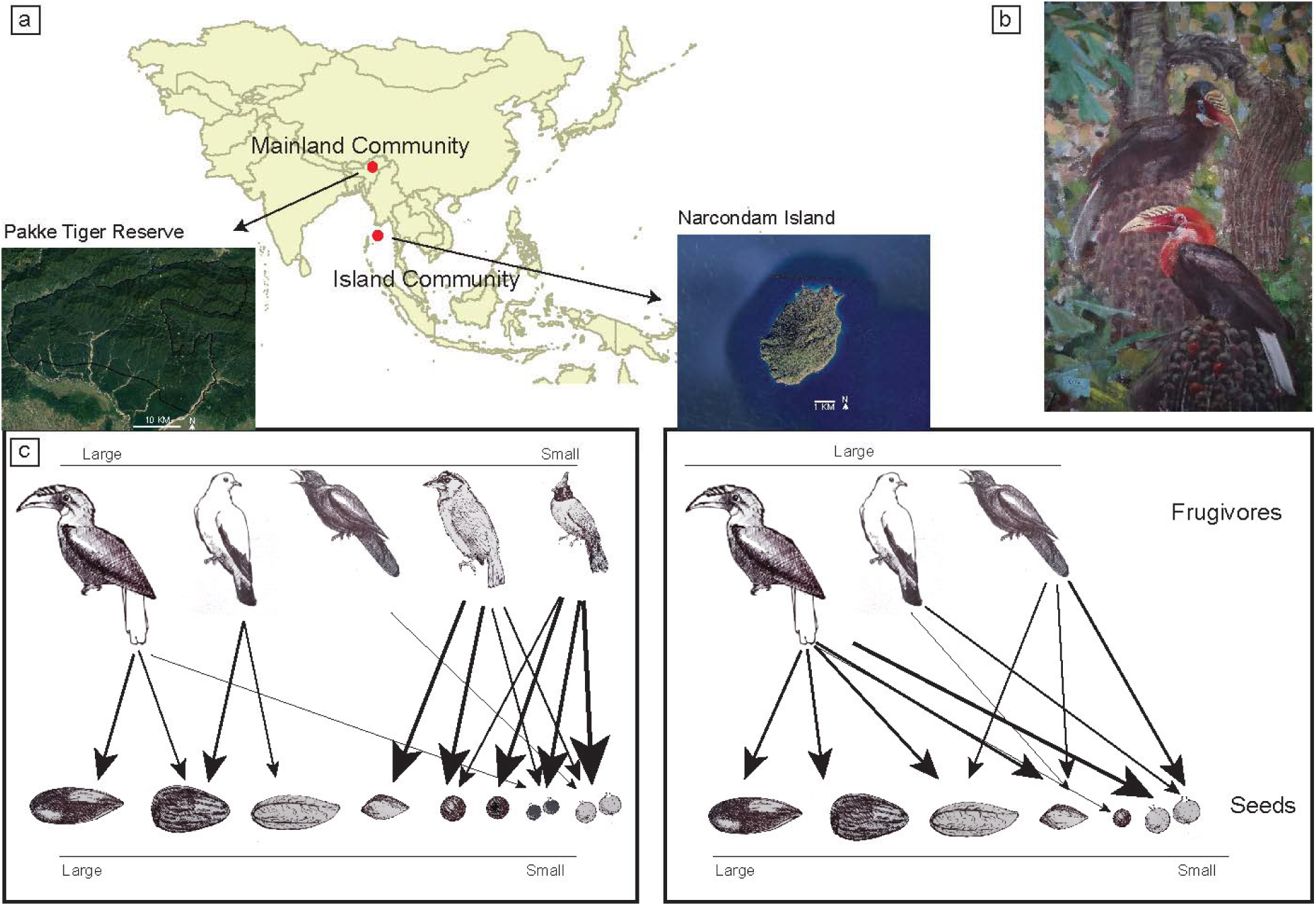
Schematic illustrating mainland and island plant and seed-dispersal communities. a) map of the South and Southeast Asia showing mainland and island site, b) the island endemic large-bodied frugivore Narcondam Hornbill *Rhyticeros narcondami* on *Caryota mitis*, c) Representative schematic showing organization of plant-seed disperser community on mainland and island. Mainland community is more diverse and symmetric as compared to the island community. On mainland, due to the presence of great diversity of small-bodied frugivores which are more efficient in removal of small fruits, large bodied frugivores may predominantly feed on large-seeded plants. On the island, in the absence of small-bodied frugivores, large-bodied frugivores will start feeding on small fruits resulting in dietary shifts and more dominant role in the community as indicated by the thickness of arrows. Dietary changes will in turn result in changes in interaction frequencies with corresponding changes in network structure. Painting and sketches by Sartaj Ghuman. Map source: Google Earth.

The influence of phylogenetic component would indicate the role of evolutionary processes like co-evolution in the organization of the community, while the influence of functional component controlling for phylogenetic history would imply trait complementarity. An integrated approach that examines the role of evolutionary processes in tandem with ecological processes thus enables an understanding of the relative roles of these factors in patterns of community assembly. However, while the substantial phylogenetic data now available has improved our understanding of community assemblages, the complementary empirical data that would allow for an integrated approach has been lacking.

Despite zoochory being a pervasive interaction in the wet tropics of Asia, there is limited information on plant-disperser communities from the mainland sites in the region (Corlett 2017) and very poor information from oceanic islands in the Indian Ocean (Escribano-Avila et al. 2018). Most of the information on plant-disperser communities worldwide comes from mainland sites with few studies from oceanic islands and even few studies set in an island-mainland comparative framework (Schleuning et al. 2014).

Given this background, we examined the organization of the plant-disperser community on a remote and small tropical island (Narcondam Island) in South Asia using the network and ecophylogenetic approaches (Fig. 1A). The Narcondam island, which is the only place where the Narcondam Hornbill *Rhyticeros narcondami* is found (Fig. 1B), is characterized by a species-poor frugivore assemblage, which also enabled us to examine the relative quantitative role of different species in seed dispersal.

In the Asian tropics, small-bodied frugivores like bulbuls and barbets are known to play key roles in seed dispersal (Naniwadekar et al. 2019). However, the Narcondam Island does not have any bulbuls and barbets and most of the dominant frugivores on the island are medium- and large-bodied avian frugivores (> 100 g) (Fig. 1C). This offers an interesting system to understand the pattern of interactions in absence of key frugivore group. We compared this island community with a mainland community from north-east India with which it shares several frugivore and plant species.

The specific aims of the study were to 1) identify key seed dispersers and plant groups on an oceanic island community by comparing species-level properties (degree, species strength and weighted betweenness) across different species, 2) examine the influence of abundance and morphological traits on species strength of plants and frugivore species, 3) compare the network-level properties (web asymmetry, connectance, specialization and weighted nestedness) of the island community with a mainland community, and 4) examine the influence of phylogenetic and functional component (fruit and seed size for plants and body size and degree of frugivory for frugivores) on the distribution of interactions between plants and frugivores on the island and mainland communities. This system allowed us to explicitly examine the change in the role of large-bodied frugivores in plant-seed disperser communities in the absence of the important small-bodied frugivores in Asian tropics (Fig. 1C). We hypothesized that large-bodied frugivores, unhindered by morphological constraints, will demonstrate dietary shifts on islands in the absence of small-bodied frugivores and will feed on fruits which they typically do not feed on in the mainland site. This will alter the organization of the plant-seed disperser community on the island as compared to mainland.

## MATERIALS AND METHODS

### Study area

The study was carried out on a small (area: 6.8 km^2^), volcanic island, the Narcondam Island (13°30’N, 94°38’E), in the Andaman Sea in India. The island is part of the Indo-Myanmar Biodiversity Hotspot (Myers et al. 2000). Three distinct vegetation types on the island include the narrow strip of littoral forests, forests dominated by deciduous species in the north-eastern part of the island and evergreen forests in most of the island (Page et al. in prep.).

The Endangered Narcondam Hornbill *Rhyticeros narcondami* is found only on the Narcondam Island (BirdLife International 2017). Around 50 species of birds have been reported from the island (Raman et al. 2013). There are two terrestrial mammals on the island, including a shrew (*Corcidura* sp.) and an invasive rat (*Rattus* sp.). There are at least four species of bats on the island including a fruit bat species (*Pteropus hypomelanus*). Narcondam Hornbill (body mass: 600-750 g) is the largest frugivore on the island. Additional details of study area are in ESM Online Resource 1.

### Frugivore and plant abundance

We conducted the study between December 2019 – February 2020. We used variable-width line-transect surveys to estimate the density and abundance of birds on Narcondam Island (Thomas et al. 2010). We walked 20 unique trails on the island (ESM Online Resource 2: Fig. S1). We divided the entire elevation gradient into three elevation zones: low (0-200 m), mid (200-400 m) and high (400-700 m) based on topography, vegetation structure and composition. One to three observers walked trails in the mornings and afternoons. During the trail walks, we recorded species identity, the number of individuals and the perpendicular distance of the sighting to the trail. The trail lengths varied between 0.22–1.26 km and the total sampling effort was 51.4 km (ESM Online Resource 2: Table S1).

We laid 49 belt transects (50 m × 10 m) across the three different elevation zones (low: 18, mid: 14, high: 17) to estimate the richness and abundance of woody plants on the Narcondam Island. Within the belt, we enumerated all wood plants that were ≥ 10 cm GBH (girth at breast height) along with their girth and taxonomic identity.

### Plant-frugivore interactions

To systematically document plant-frugivore interactions, we performed spot-sampling during our trail walks, and opportunistically outside our trail walks following Palacio *et al*. (2016). Whenever we encountered a frugivore on a fruiting tree, we recorded the species identity, number of frugivores, and frugivore foraging behaviour (fruits swallowed, pecked or dropped) to ascertain whether the frugivore was a seed disperser or not. Plant-frugivore interaction information was systematically collected outside trail sampling whenever frugivore was detected on fruiting trees. To determine if we were missing documenting interactions of frugivore species with low detection probability through spot sampling, we systematically observed fruiting trees. We observed 40 fruiting individuals of 12 species following Naniwadekar et al. (2019a) (ESM Online Resource 2: Table S2). Additional details of the tree watches are in ESM Online Resource 1. We measured widths of at least five fruits each of 23 species of fleshy-fruited plants following Naniwadekar et al. (2019*a*) (ESM Online Resource 2: Table S3). Plants were classified as small-seeded (seed width: < 5 mm), medium-seeded (seed width: 5-15 mm) and large-seeded (> 15 mm) following Naniwadekar et al. (2019a) (ESM Online Resource 2: Table S3).

### Analysis

#### Frugivore and plant abundance

We estimated densities of perched birds with more than 40 visual detections using the R package ‘Distance’ (Thomas et al. 2010; Miller et al. 2019). We estimated combined density of two Imperial*-*pigeon species (Pied Imperial-Pigeon *Ducula bicolor* and Green Imperial-Pigeon *Ducula aenea*), since we had 45 detections overall. Additional details of the analysis are provided in the ESM Online Resource 1. We estimated the overall tree density and basal area (m^2^ ha^-1^), and density of 27 species of fleshy-fruited plants that were fruiting during the study period.

#### Island plant-seed disperser community

We created a matrix of plants and frugivores with the frequency of interactions documented for each plant-frugivore combination through spot sampling. We only considered those animals as seed dispersers which we had seen swallowing the seeds. We didn’t consider Alexandrine Parakeet *Psittacula eupatria* in the analysis as we documented the parakeets predating on seeds or feeding on unripe fruits as has also been documented elsewhere (Shanahan et al. 2001). We plotted the interaction accumulation curve using the ‘random’ method and 1000 permutations as implemented in the R package ‘vegan’ to determine if we had adequately documented the diversity of unique plant-seed disperser interactions (Oksanen et al. 2019).

To determine the relative roles of species in the organization of plant-seed disperser communities, we examined species-level properties like the degree, species strength and weighted betweenness centrality. Degree is defined as number of mutualistic partners of the focal species, species strength is the sum of level of dependencies of all the partner species on the focal species and betweenness centrality is a useful measure to identify connector species which connects multiple guilds in a community (Dormann et al. 2008; Martín González et al. 2010). We used the R package ‘bipartite’ for this analysis (Dormann et al. 2008).

We used the general linear model to examine the influence of plant abundance (estimated using the vegetation plots) and fruit type (small-seeded, figs, medium-seeded and large-seeded) on species strength of the 27 plant species that were fruiting during the study period. During exploratory analysis, we noticed that figs consistently had higher species strength values. Since certain fig groups are known to be keystone species in tropical forests and provide key nutrients (Shanahan et al. 2001), we performed the analysis with figs as a separate category. Species strength was log_10_ transformed to approximate normality. In the case of frugivores, since there were only seven species and species strength values were not normally distributed, we used Spearman’s *ρ* to examine the correlation between abundance (encounter rate km^-1^) of frugivores and body mass with species strength. We obtained body mass information from Wilman et al. (2014). To determine if the island plant-seed disperser community was organized into distinct sub-communities, we used the program MODULAR (Marquitti et al. 2014) following Naniwadekar *et al*. (2019). Additional details in ESM Online Resource 1.

#### Comparing the island and the mainland plant-seed disperser communities

To determine the differences in the organization of plant-seed disperser communities between island and mainland, we compared plant-disperser communities between the Narcondam Island with that of Pakke Tiger Reserve in north-east India. Information on plant-seed disperser communities is limited from the Asian tropics. However, there is detailed information on interactions between 47 avian frugivores and 43 plant species is available from Pakke Tiger Reserve in north-east India (Naniwadekar et al. 2019). Pakke is part of the Eastern Himalaya Biodiversity Hotspot. It is about 1500 km straight line distance north from the Narcondam Island. Unlike the island community which is relatively young, the mainland site is several million years old.

Representative species of all the frugivore genera found on the Narcondam Island are also found in Pakke with species like the Green Imperial-Pigeon, Asian Koel *Eudynamys scolopaceus* and Common Hill Myna *Gracula religiosa* being reported from both sites. Several genera of fleshy-fruited plants found on Narcondam Island are also found in Pakke. We examined network-level properties like web asymmetry, connectance (proportion of realized interactions), specialization (*H*_*2*_*’*) and nestedness (weighted NODF) on Narcondam Island and compared it with the mainland site (Pakke). Values of web asymmetry range between −1 to 1 with negative values indicating higher diversity of plant species as compared to animal species and vice versa (Blüthgen et al. 2007). Specialization is a measure of complementary specialization of the entire network with high values indicating that the species are more selective. Weighted NODF is a quantitative measure of nestedness with high values indicating higher nestedness.

#### Community assembly on island and mainland

To determine the influence of phylogeny and functional traits on plant-frugivore interactions on the mainland and island site, we used the following approach. The plant phylogeny was pruned from the megaphylogeny (Qian and Jin 2016) for both the island and mainland communities individually. This is a time-calibrated phylogeny with largest representation from 98.6% of families and 51.6% of all genera of seed plants. Scientific names of plants were standardized by referring to ‘The Plant List’ (www.theplantlist.org). However, some species from Pakke and Narcondam were missing from the megaphylogeny. In those cases, we chose the closest species from the Oriental region. When we failed to find a congener, we chose a genus close to the focal genus (ESM Online Resource 2: Table S3). We used fruit width and seed size class information for the analysis (ESM Online Resource 2: Table S3). In the case of the birds, we referred to the comprehensive and time-calibrated bird phylogeny available at www.birdtree.org (Jetz et al. 2012). We found all the focal species for the two sites in the database except *Pycnonotus flaviventris, Chrysocolaptes guttacristatus* and *Sitta cinnamoventris*. We used congeners from the Oriental region for these species for the analysis (ESM Online Resource 2: Table S4). For the frugivores, we used body mass and percentage fruits in the diet of frugivores (degree of frugivory) obtained from Wilman et al. (2014) (ESM Online Resource 2: Table S4). We obtained 100 trees from posterior distribution of 10000 trees based on Ericson Sequenced Species backbone phylogeny with 6670 OTUs for seven bird species for Narcondam and 47 species for Pakke found in the network (Jetz et al. 2012). The maximum clade credibility tree was used for further analyses as these trees did not vary substantially in topology and branch lengths. We calculated the phylogenetic distances (pair-wise distances between species using their branch lengths) using ‘cophenetic.phylo’ in the R package ‘ape’ (Paradis and Schliep 2018). We used Gower method to estimate pair-wise distances among traits of both plants and frugivores using “vegdist” in the R package ‘vegan’. We then followed the novel approach outlined by Bastazini et al. (2017) to determine whether phylogenetic proximity influenced the evolution of species trait and their interactions in the network or whether species traits influence the observed pattern of interactions after removing the potential confounding effect of the phylogeny. We used a weighted matrix of interactions between plants and frugivores. It was frequency of interactions for the island site and visitation rates for the mainland site. We applied this method for Narcondam and Pakke separately. We used the R packages ‘SYNCSA’ for this analysis (Debastiani and Pillar 2012; Oksanen et al. 2019). We used R (ver. 3.5.3) for all the analysis (R Core Team 2019).

In addition, we assessed the phylogenetic signal separately in trait data for birds and plants in both sites, Narcondam and Pakke. We calculated the Blomberg’s *K* for each trait using the R package “phytools” (Blomberg et al. 2003; Revell 2012).

## RESULTS

### Frugivore and plant abundance on island

We detected six avian frugivores during the trail walks. We had 207 visual detections of Narcondam Hornbill, 71 detections of Asian Koel, and 45 detections of the two Imperial-Pigeons. We had only two detections each of the Common Hill Myna and Eye-browed Thrush *Turdus obscurus*. Information on detection probability and mean flock size is summarized in ESM Online Resource 2: Table S5. Narcondam Hornbill was the most common frugivore on the island and its abundance (mean (95% CI): 1026 birds (751-1402)) was almost four times that of the Asian Koel (Fig. 2A; ESM Online Resource 1: Table S5). The estimated overall mean (95% CI) density of the Narcondam Hornbill was 151 (110-206) birds km^-2^. The mean densities of the Narcondam Hornbill varied between 101-168 birds km^-2^ across the three elevation zones with the lowest mean densities in the highest elevation band (Fig. 2B) (ESM Online Resource 2: Table S5).

**Figure 2.**
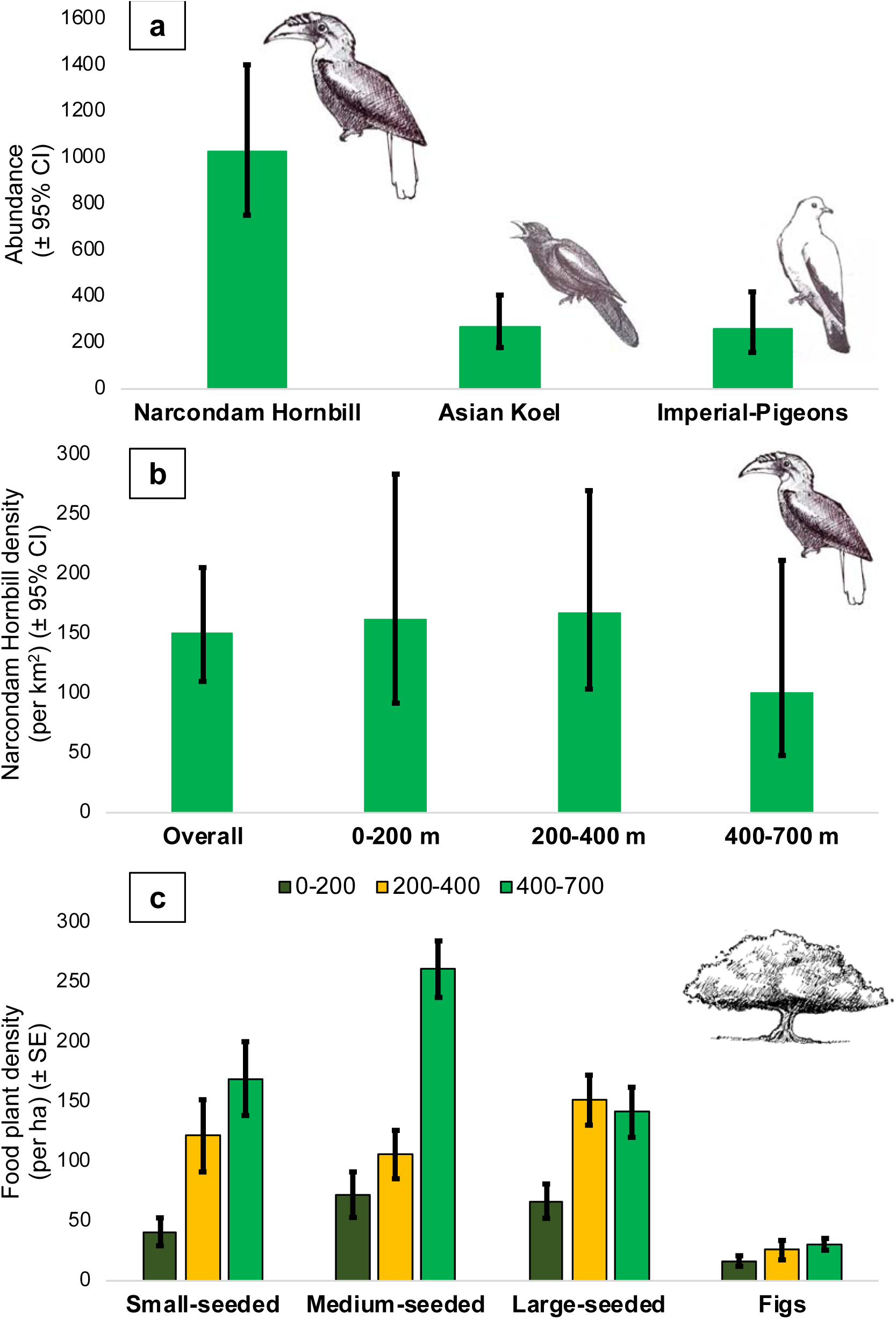
Abundance of Narcondam Hornbill *Rhyticeros narcondami*, Asian Koel *Eudynamys scolopaceus* and Imperial-Pigeons (Pied Imperial-Pigeon *Ducula bicolor* and Green Imperial-Pigeon *Ducula aenea*) on Narcondam Island (A). Density (km^-1^) of Narcondam Hornbill on the island and across the three elevation zones on the island (B). Density (ha^-1^) of 27 species of fleshy-fruited plants that were fruiting across the three elevation zones during the study period (C).

The mean (± SE) density and basal area of all the woody plants (≥ 10 cm GBH) was highest in the 0-200 m elevation band (density: 2091.1 ± 124.4 plants ha^-1^; basal area (m^2^ ha^-1^): 58.3 ± 5.3), followed by 200-400 m elevation band (density: 1505.7 ± 101.1 plants ha^-1^; basal area (m^2^ ha^-1^): 49.1 ± 3.0) and least in the 400-700 m elevation band (density: 1311.8 ± 82.7 plants ha^-1^; basal area (m^2^ ha^-1^): 39.2 ± 4.2). However, the mean abundance of the different categories of the frugivore food plant species (figs, small-seeded non-fig plants, medium-seeded plants and large-seeded plants), which were fruiting during the study period, was highest in the high elevation band (400-700 m), followed by middle elevation band and lowest in the low elevation band (0-200 m) (Fig. 2C).

### Island plant-seed disperser community

We documented 752 interactions between seven frugivore species and 27 plant species during the spot scans performed on trail walks and opportunistically. We documented all the frugivore species (except Andaman Green Pigeon *Treron chloropterus*) previously reported from the island. We documented the migrant Daurian Starling *Agropsar sturninus*, for the first time. During spot scans on trail walks, we documented 262 interactions between six frugivore species and 19 plant species. During opportunistic spot scans, we documented 490 interactions between seven frugivore species and 26 plant species. Narcondam Hornbill was documented feeding on 22 plant species, followed by Asian Koel, which was documented feeding on 16 plant species (ESM Online Resource 2: Table S6). Of the 752 interactions, 533 were of the Narcondam Hornbill, 111 were of the Asian Koel, 36 of the Green Imperial-Pigeon, 34 of the Pied Imperial Pigeon, 25 of the Common Hill Myna, 12 of the Eye-browed Thrush and one of the Daurian Starling. Among the 27 plant species, eight species were represented by *Ficus*. We observed 282 interactions on *Ficus rumphii*, 80 on *Ficus glaberrima*, 54 on the small-seeded *Aidia densiflora* and 50 on the large-seeded *Caryota mitis*. We documented 64 unique interactions between plants and frugivores on the island during the study period. The interaction accumulation curve indicated that we had adequately documented interaction diversity (ESM Online Resource 2: Fig. S2).

During tree watches, we documented 684 interactions (Narcondam Hornbill: 481, Asian Koel: 167, Pied Imperial-Pigeon: 23, Green Imperial-Pigeon: 7, Eye-browed Thrush: 5, Common Hill Myna: 1), between 11 plant species and six frugivore species. We did not document any interaction during tree watches which we had not documented during the spot scans. For eight of the 11 tree species, we observed greater diversity of frugivores foraging on the particular plant species during spot sampling as compared to tree watches.

*Ficus rumphii* (among plants) and Narcondam Hornbill (among frugivores) had the highest degree, species strength and weighted betweenness centrality highlighting the central role they played in the community during the study period (ESM Online Resource 2: Table S6). General linear model indicated that the species strength in plants was influenced by plant type (figs and non-fig small-, medium- and large-seeded plants) and not by plant abundance (Adj. *R*^*2*^ = 0.40, *F*_4,22_ = 5.322, *p* = 0.004) (Fig. 3A; ESM Online Resource 2: Table S7). *Ficus* were different from other small-seeded plants and had higher species strength values (mean (± SE) = 0.739 (± 0.101), n = 8 species) as compared to small-seeded (mean (± SE) = 0.0816 (± 0.01), n = 9 species), medium-seeded (mean (± SE) = 0.0376 (± 0.01), n = 6 species) and large-seeded plant species (mean (± SE) = 0.0328 (± 0.01); n = 4 species) (Fig. 3A). Among frugivores, species strength was significantly and positively correlated to their abundance (Spearman’s *ρ* = 0.89, *p* = 0.0123, n = 7 species; Fig. 3B) and not significantly correlated to their body mass (Spearman’s *ρ* = 0.71, *p* = 0.088, n= 7).

**Figure 3.**
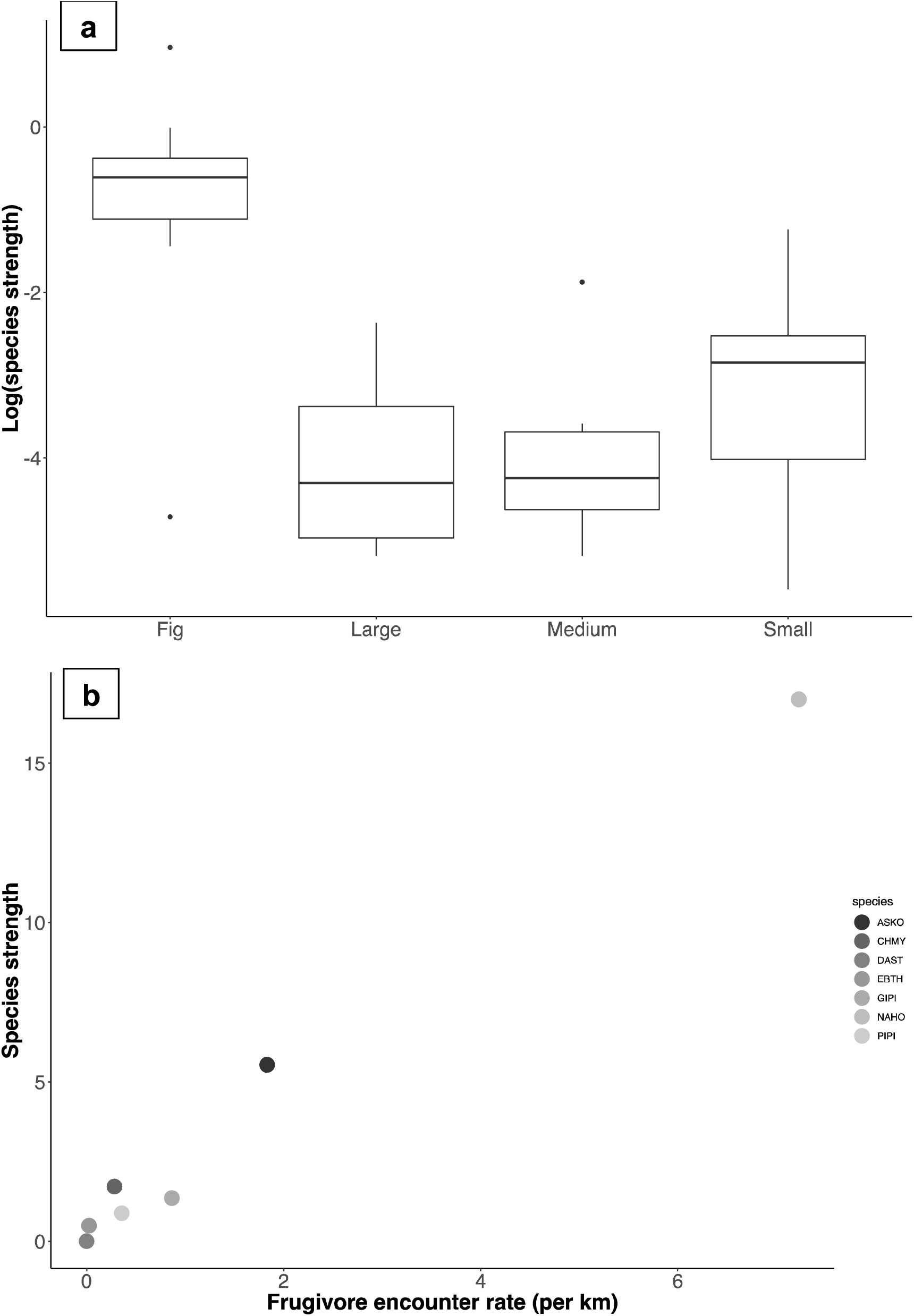
Relationship between species strength (log_10_ transformed in the case of plants) and plant type (A) and frugivore abundance (B). We separated figs and small-seeded plants for they consistently had higher species strength values. Frugivore codes - ASKO: Asian Koel, CHMY: Common Hill Myna, DAST: Daurian Starling, EBTH: Eye-browed Thrush, GIPI: Green Imperial-Pigeon, NAHO: Narcondam Hornbill and PIPI: Pied Imperial-Pigeon.

The observed modularity of the island plant-disperser community was 0.33, and it was not significantly different from that obtained through the null models (*p* = 0.45). The observed community was subdivided into four modules and all the species were consistently allocated to the same modules across the 25 runs (Table S8). One of the four modules comprised of the Narcondam Hornbill and several medium and all the four large-seeded plants including *Endocomia macrocomia, Caryota mitis, Pycnarrhaena lucida*, and *Planchonella longipetiolata* highlighting hornbill’s importance for large-seeded plants (Table S8).

### Comparing the island and the mainland plant-seed disperser communities

The island network was more asymmetric (−0.588) as compared to the mainland site (0.044). The observed values of connectance and nestedness on Narcondam were almost twice that of the mainland site and specialization (*H*_*2*_*’*) was almost half that of the mainland site (Table 1). The observed network on Narcondam was less connected and less weighted than the null model predictions but it was more specialized than expected by random (Table 1).

**Table 1.** Network-level properties (observed and null model values) for the Narcondam Island, and observed values for the mainland site, Pakke Tiger Reserve. Null model values for the mainland site can be found in Naniwadekar et al. (2019a).

|  | **Observed value:**<br>**Narcondam** | **Null mean:**<br>**Narcondam** | **Null SD:**<br>**Narcondam** | **Observed value:**<br>**Mainland** |
| --- | --- | --- | --- | --- |
| Connectance | 0.339 | 0.463 | 0.018 | 0.154 |
| Specialization ($H_2'$) | 0.233 | 0.144 | 0.009 | 0.500 |
| Weighted NODF | 45.178 | 59.76 | 3.834 | 24.885 |

### Community assembly on island and mainland

Among woody plants, we detected a strong phylogenetic signal for seed size and fruit width in Pakke Tiger Reserve (seed size: Blomberg’s *K* = 0.73, *p* = 0.001; fruit width: Blomberg’s *K* = 0.43, *p* = 0.005) and Narcondam (seed size: Blomberg’s *K* = 1.48, *p* = 0.001; fruit width: Blomberg’s *K* = 0.45, *p* = 0.027). Among frugivores, we detected a phylogenetic signal for the body mass of frugivores in Pakke (Blomberg’s *K* = 1.22, *p* = 0.012) and Narcondam Island (Blomberg’s *K* = 1.28, *p* = 0.043). We did not detect phylogenetic signal in the percentage of fruits in the diet of the frugivores of Pakke (Blomberg’s *K* = 0.23, *p* = 0.38) and Narcondam (Blomberg’s *K* = 0.90, *p* = 0.08). We did not find any significant correlation for the four models indicating that neither the functional nor the phylogenetic component was able to explain the observed interaction patterns between plants and frugivores in both island and mainland sites (Table 2; Fig. 4).

**Table 2.**
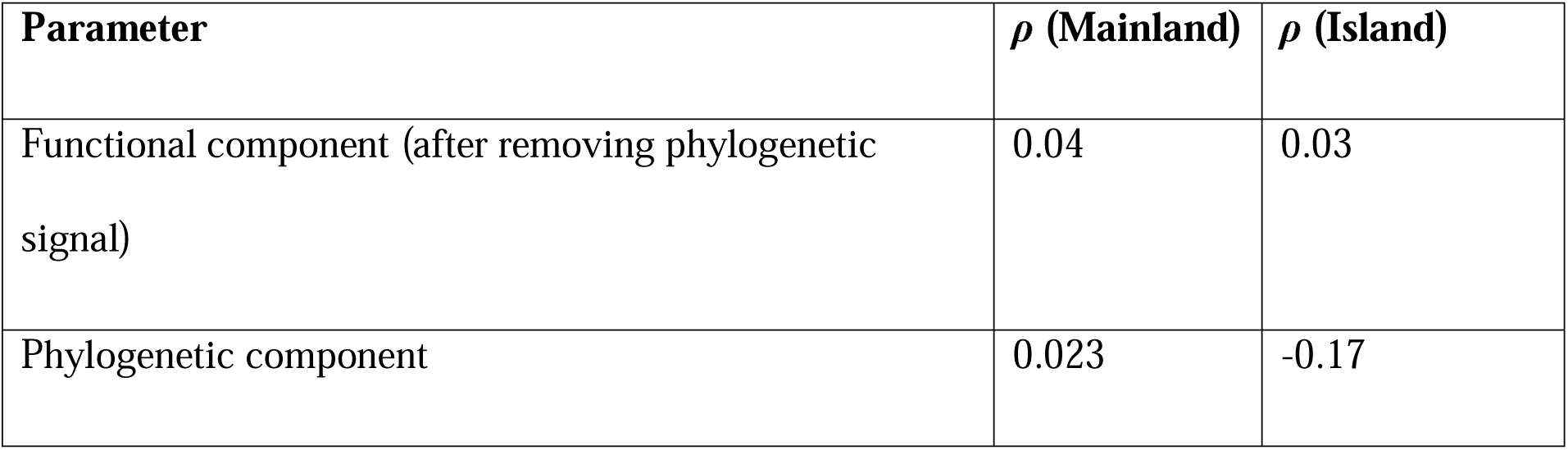
Correlation coefficients for the two models that evaluate the influence of the functional (after removing phylogenetic signal) and phylogenetic components on the observed patterns of interactions between plants and frugivores for the mainland site (Pakke Tiger Reserve) and the island site (Narcondam Wildlife Sanctuary). None of the values were significant at *p* = 0.05.

| Parameter | $\rho$ (Mainland) | $\rho$ (Island) |
| --- | --- | --- |
| Functional component (after removing phylogenetic signal) | 0.04 | 0.03 |
| Phylogenetic component | 0.023 | -0.17 |

**Figure 4.**
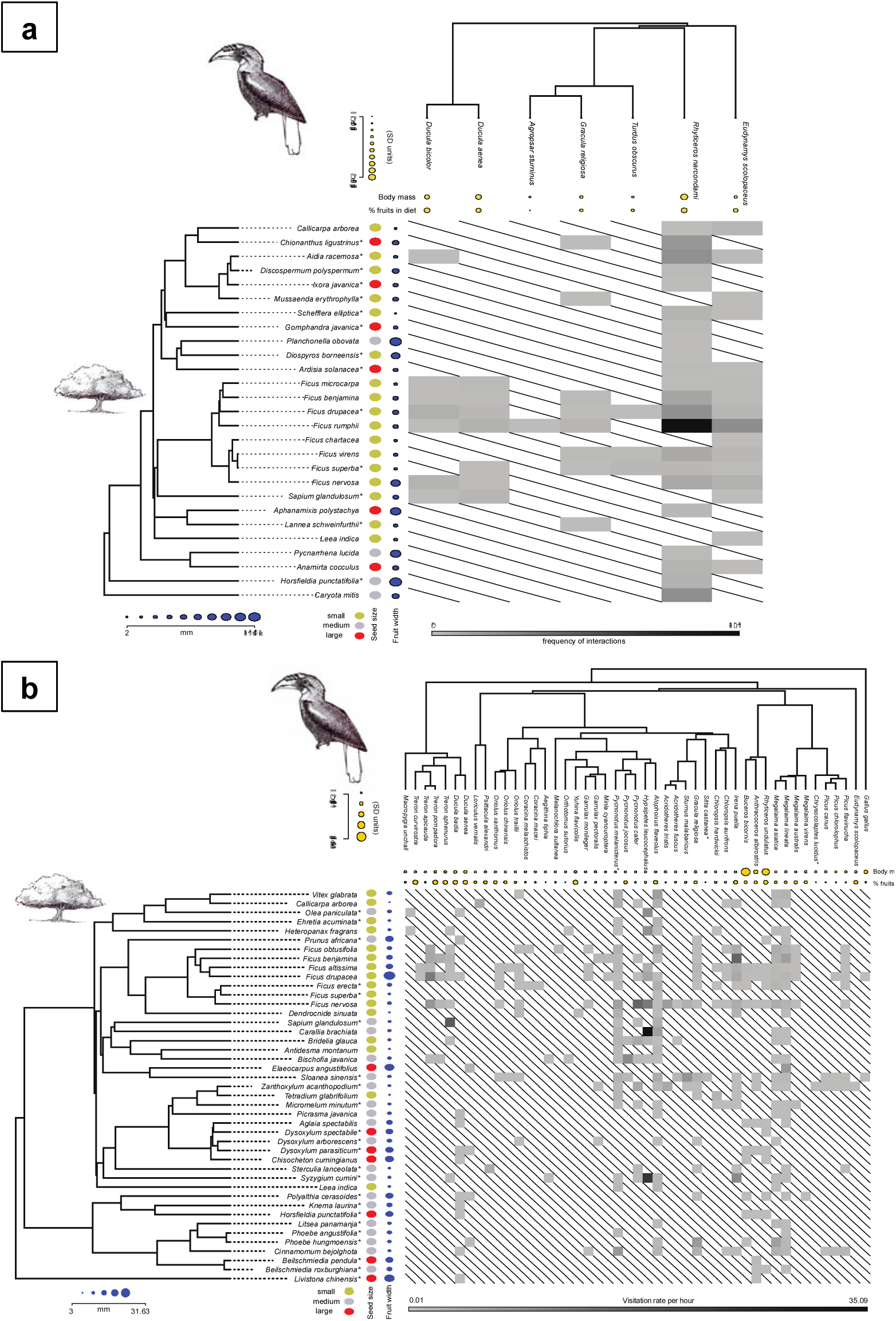
Seed dispersal interactions in Narcondam Island (A) and the mainland site, Pakke Tiger Reserve (B). Also shown is the pruned phylogeny for the plants and the frugivores and respective traits for plants (seed size and fruit width) and frugivores (body mass and percentage diet that is comprised of fruits). The matrix is weighted in both the island and mainland case and it has frequency of interactions for the island site and visitation rate of frugivores (h^-1^) for the mainland site shown by the grey bar below the matrix. * indicates species for which a congener or allied genus was used in the constructing the phylogenetic tree.

## DISCUSSION

In this study, we demonstrate the role of large-bodied frugivores, especially the endemic Narcondam Hornbill, in contributing disproportionately to seed dispersal in the absence of key small-bodied frugivore groups on the island. Narcondam Hornbill plays a central role in the plant-dispersal community as it is super-abundant, has very high species strength and betweenness centrality values and is critical for the select large-seeded plants fruiting. This study also demonstrates the key role played by figs in the plant-seed disperser community as demonstrated by its very high species strength values as compared to other plant types. In comparison to the mainland community, the island community was more asymmetric, connected, nested and less-specialized. We have documented trait convergence and dietary shifts in frugivores which potentially contribute to the diffused nature of plant-seed disperser interactions. This also likely contributes to the absence of the role of phylogenetic or functional component in the assembly of plant-seed disperser communities on mainland or the island. This is one of the few studies from oceanic islands of the Indian Ocean and demonstrates how the largest frugivore, which is a point-endemic, plays a critical quantitative role in seed dispersal. This is also one of the few studies that performed a paired comparison between an island and mainland community.

### Narcondam for the hornbill and by the hornbill

There is no other place in the world where hornbills are reported to occur in such high densities as on Narcondam Island (151 birds km^-2^). On Sulawesi, the Knobbed Hornbills *Rhyticeros cassidix* occur in very high densities of up to 84 birds km^-2^ (Kinnaird et al. 1996). In north-east India, Wreathed Hornbill *Rhyticeros undulatus*, a close relative of the Narcondam Hornbill, seasonally occur in densities of up to 68 birds km^-2^ (Naniwadekar and Datta 2013). Our density estimates are similar to densities estimated by previous studies (Raman et al. 2013; Manchi 2017). Reduction in hunting pressures may have resulted in the recovery of Narcondam Hornbill population from around 300 – 400 birds in early 2000s following the suggestions by Sankaran (2000). Among other frugivores, there is limited information on the density of the Asian Koel from other sites, but comparison with existing studies show that Narcondam has very high densities of as compared to other sites (Riley 2001). The mean Imperial-Pigeon densities are slightly lower than those reported from a mainland site in north-east India (Dasgupta and Hilaluddin 2012).

In this study, multiple measures (abundance and different network metrics – species strength, degree, betweenness and modularity) indicate that Narcondam Hornbills are keystone seed dispersers on the island. Generally, as the body mass increases, the abundance of the frugivore and the frequency of visitation is found to decline (Godínez-Alvarez et al. 2020). Interestingly, the Narcondam Island is unique where the largest frugivore, the Narcondam Hornbill, is also the most abundant frugivore on the island. Abundant species have been reported to influence their partners strongly and Narcondam Hornbill had the highest species strength values highlighting its importance for plants (Vázquez et al. 2007). Larger frugivores, like hornbills, are known to remove more fruits per visit as compared to other smaller frugivores (Naniwadekar et al. 2019). Highest number of interactions of the Narcondam Hornbill coupled with potentially higher fruit removal rates point towards the key quantitative role played by the hornbill on the island.

Modularity analysis revealed that the hornbill was the key seed disperser for the large-seeded plants as has been demonstrated in previous studies (Naniwadekar et al. 2019; Gopal et al. 2020). The hornbill was the only frugivore that has high betweenness centrality value pointing towards its vital role as a connector between different guilds within the community. This high reliance on a frugivore and the potentially low redundancy highlights the fragility of the island ecosystem because of strong reliance on a single frugivore.

Narcondam has very high densities of hornbill food plants, which likely explains the high hornbill densities. The overall fig density on Narcondam Island was 27 trees ha^-1^. This is very high as compared to other sites in the world where the densities of hemi-epiphytic figs are typically less than one tree ha^-1^ (Harrison 2005). *Ficus* (overall) densities were 1.4 trees ha^-1^ at a site in the southern Western Ghats (Page and Shanker 2020). *Ficus* densities of up to 10 trees ha^-1^ have been reported in Sulawesi (Kinnaird et al. 1996). Consistently across the mainland site and the island site, certain species of *Ficus* attracted the highest diversity of frugivores. On the island, *Ficus* represented the top six species in species strength, highlighting the key role they potentially played in the plant-disperser community at least during the study period. Association between canopy, hemi-epiphytic figs and hornbills has been documented elsewhere (Harrison et al. 2003). Most dominant fig species during our study on Narcondam Island, which are important hornbill food plants are mostly canopy, hemi-epiphytic figs. Being small-seeded, they often play the pioneering role in attracting frugivores on oceanic islands and accelerating plant colonization on the island (Harrison 2005). Figs and hornbills may have played a critical role in setting up the plant community on the island which needs further investigation.

Based on published literature on hornbill food plants from north-east (Datta 2001; Naniwadekar et al. 2015, 2019) and Thailand (Poonswad et al. 1998; Kanwatanakid-Savini et al. 2009; Chaisuriyanun et al. 2011), we identified at least 35 species of Narcondam Hornbill food plants. Overall density of the 35 hornbill food plant species (GBH ≥ 30 cm) on Narcondam Island is 327 trees ha^-1^. In Pakke Tiger Reserve, the density of 45 hornbill food plant species was 163 trees ha^-1^ (Datta 2001), highlighting the hyper abundance of hornbill food plants on Narcondam Island. Interestingly, the abundance of food plants is consistently distributed across the entire elevation gradient. The hyper abundance of frugivores and their food plants makes this island a unique ecosystem.

### Plant-seed disperser communities on island and mainland

Island communities have been reported to be more asymmetric as compared to mainland systems due to higher immigration rates of plants or human-caused frugivore extinctions on the island (Schleuning et al. 2014; Nogales et al. 2016). There are no known human-driven extinctions on the Narcondam island. We are unlikely to find additional species of frugivores that can play an important functional role in seed dispersal on the island as our list of frugivores is comparable to the frugivores reported by the previous studies (Raman et al. 2013). We feel that dispersal limitation of frugivores could also be a key factor in the web asymmetry. Andaman and Nicobar archipelago of which Narcondam Island is part of, is an interesting example of this. Bulbuls and barbets, which are the key frugivores on the mainland (Naniwadekar et al. 2019), are absent on the remote and tiny Narcondam Island. Endemic species of bulbuls are found on other islands in the Andaman and Nicobar archipelago, but barbets are absent from the archipelago. Thus, the variation in the dispersal ability of different frugivore groups might also play an important role in contributing towards the web asymmetry on oceanic islands. Among other network properties, higher connectance on islands has been reported elsewhere (González-Castro et al. 2012).

Forbidden links are thought to contribute towards lower connectance (Olesen et al. 2011). Additionally, connectance and nestedness are negatively correlated (Rezende et al. 2007). In our case, higher connectance is associated with higher nestedness on the island site as compared to the mainland site. Given that most of the important frugivores on the island are medium (100-500 g) or large-bodied (> 500 g) with ability to handle small and large fruits (thereby less likelihood of forbidden links), it has likely resulted in higher connectance and nestedness.

### Organization of plant-seed disperser communities

We detected contrasting phylogenetic signals in bird body mass, and seed and fruit traits on both the island and the mainland. While frugivore traits were more likely to be similar (than Brownian motion model) for closely related species, plant traits were less likely to be similar for closely related species. However, functional and phylogenetic component did not influence the network interactions in the plant-seed disperser communities on the island and mainland. The role of evolutionary history in shaping local networks of interacting species is thought to be limited (Segar et al. 2020). Unlike the plant-pollinator networks, the seed dispersal networks can be expected to be less specialized and diffused as unrelated species may exhibit trait convergence and forage on similar plant species (Jordano 2000). We found that distantly related groups of frugivores dispersed seeds of a common assemblage of plants, many of which are distantly related to each other. A diverse array of frugivores disperse the super-generalist *Ficus*, and even the large-seeded plants, which otherwise have a smaller disperser assemblage, were predominantly dispersed by distant frugivore clades, like hornbills and *Ducula* pigeons, on the mainland site. Unlike Bastazini et al. (2017), we did not detect signal in the functional component of the plant-seed disperser communities. This is a likely consequence of the absence of trait complementarity among interacting frugivores and plants. While the small-bodied frugivores may not be able to disperse large seeds due to morphological constraints, large-bodied frugivores like hornbills, fed on plants that were large- and small-seeded resulting in lack of trait complementarity.

Another important reason could be the behavioral plasticity shown by frugivorous birds while foraging. In the mainland site, where there are almost 50 frugivore species with a higher preponderance of small-bodied frugivores, we rarely encountered *Ducula* pigeons on figs that attracted more than half of the frugivore species found in the area. On the mainland site, *Ducula* pigeons were mostly observed foraging on medium-seeded and large-seeded plants. On the island, where there are hardly any small frugivores, we mostly saw the *Ducula* pigeons foraging on *Ficus*. This behavior of *Ducula* pigeons feeding on figs on the island but less frequently on the mainland demonstrates phenotypic plasticity in *Ducula* pigeons, likely as a response to lack of competitive interactions, an aspect that has been seldom documented elsewhere. Additionally, on Narcondam Island, we documented hornbills feeding on *Callicarpa arborea*, a small-seeded species which is mostly dispersed by bulbuls and barbets on the mainland site (Naniwadekar et al. 2019). Asian Koel, which is a rare frugivore in the mainland site, was seen only once feeding on fruit there. However, on the island, it is the second-most important frugivore. It remains to be determined, whether exploitative competition on the mainland site drives larger frugivores to feed on different resources in the mainland forested system. While habitat fragmentation often results in loss of large-bodied frugivores thereby affecting large-seeded plants, it will be interesting to investigate the implications of the absence of small-bodied frugivore on the structuring of plant communities.

Bats are known to play an important role in seed dispersal on oceanic islands (McConkey and Drake 2015). We recorded spat out wads of pulp of three *Ficus* spp., *Balakata*, and *Planchonella*. Direct detections of foraging bats on fruiting *Manilkara zapota* (planted), *Terminalia catappa, Ficus rumphii* and *Ardisia*. We failed to detect foraging bats during the six night-walks that lasted for at least 90 minutes each.

One may argue about the lack of phenological completeness of the study and its influence on the inferences drawn. The abundances of the frugivores is unlikely to change since it is a small and remote island in the middle of the Andaman Sea. Narcondam Hornbill is likely to remain the most abundant frugivore on the island year-round and thereby play a key role in seed dispersal throughout the year. Additionally, the frugivore species composition on the island is unlikely to change as previous studies have documented similar assemblage of frugivores. We have a diverse representation of plant families in our study with varying fruit (figs, drupes and arillate dehiscent capsular fruits) and seed traits thereby capturing the functional diversity in traits of fruits and seeds making the data amenable for ecophylogenetic analysis. Additionally, this is a remote island making it extremely challenging to conduct long bouts of fieldwork, thereby posing challenges to collect long-term data.

Only those plants and animals that are able to cross the oceanic barriers and establish contribute to structuring of communities on islands. Degree of isolation, area and age of islands thereby offers an interesting gradient to understand the relative importance of processes in structuring of communities. While, we did not detect a phylogenetic or functional signal across mainland and island communities, it will be interesting to investigate these patterns as the diversity of frugivores increases further. Stronger signals can be expected in a niche packing scenario. As demonstrated in this study, large-bodied frugivores, particularly the Narcondam Hornbill, play a pivotal role on Narcondam Island in the absence of small frugivores on a relatively young and moderately isolated island. Several threatened hornbill species are found on oceanic islands (IUCN 2019). Narcondam Island and its endemic hornbill give insights into the potential role of hornbills, which are among the largest frugivores on islands, in shaping and potentially maintaining the island tree communities by playing a key role as a seed disperser. The ecological role of hornbills and their contribution to mutualism and maintenance of plant species diversity is poorly understood from other oceanic islands. Narcondam Island through the hyper abundance of hornbills and its food plants and the asymmetric nature of the plant-disperser community with heavy reliance on a single frugivore demonstrates the unique ecological and evolutionary setting of oceanic islands. The disproportionate and singular role that species can play on oceanic islands points towards the extreme vulnerability of oceanic islands to any external perturbations. The perturbations can disrupt the delicate and intricate relationships between plants and seed dispersers, with dramatic cascading impacts on the entire community. Unfortunately, studies like this might not be possible in many other oceanic islands that harbor hornbills as they have been already modified extensively due to anthropogenic activities. It is indeed fortunate that Narcondam Island has been preserved as a Wildlife Sanctuary thereby safeguarding not only a unique hornbill species but also a unique ecosystem.

## Supporting information

ESM Online Resource 1

ESM Online Resource 2

## ACKNOWLEDGEMENTS

We thank Wildlife Conservation Trust, IDEAWILD, Nature Conservation Foundation, Mr. Uday Kumar, MMMRF, Mr. Rohit and Deepa Sobti, and Mr. Arvind Datar for providing funding support.We are grateful to the Andaman and Nicobar Forest Department for giving us the permit (No: WII/NVP/NARCONDAM/2019). We thank Mr. Dependra Pathak, DGP (A&N) for giving us the necessary permissions. We thank Commandant A. K. Bhama and Captain Kundan Singh from the Indian Coast Guard for giving us permission and support. We thank Mr. Abhishek Dey, DC (South Andamans) for giving us permission. We thank Kulbhushansingh Suryawanshi, Divya Mudappa and T. R. Shankar Raman for providing us field equipment and for valuable discussions. We thank Mr. D. M. Shukla (PCCF, Wildlife), Mr. A. K. Paul and Mr. Soundra Pandian for their support. We are grateful to the Dean and the then Director WII, Dr. G. S. Rawat for facilitating the research permit application and supporting the project. We are indebted to the Special Armed Police unit led by Ms. Usha Rangnani (SP) for providing us logistic support at Narcondam Island. We thank Elrika D’Souza, Evan Nazareth, Rachana Rao and Rohan Arthur for providing us logistic support in Port Blair. We thank Prasenjeet Yadav, Adarsh Raju, Suri Venkatachalam, Shashank Dalvi, Anand Osuri, Narayan Sharma and Aparajita Datta for valuable discussions. We thank Hari Sridhar, Risa Sargent and Jennifer Lau for valuable comments on a previous version of this manuscript.

## Conflicts of interest

Authors declare that they have no conflict of interest.

## Notes

### Competing Interest Statement

The authors have declared no competing interest.

### Summary of Updates

The title has been changed to better reflect the results of the study. The abstract has been edited for clarity. The introduction section has been substantially edited for clarity following feedback.

