## Supplementary material for "Large frugivores matter more on an island: insights from island-mainland comparison of plant-frugivore communities": ESM Online Resource 1

**ELECTRONIC SUPPLEMENTARY MATERIAL 1**

**Study Area:**

In the early 2000s, the Narcondam Hornbill abundances were reported to be less than 400 birds (Sankaran 2000), but a field study conducted in 2013 reported an abundance of 1293 birds (Manchi 2017). Among these studies, only one study used the conventional distance sampling approach to estimate hornbill densities and estimated a density of 167 hornbills per km^2^ in the low elevation forests (Raman et al. 2013). Previous studies did not estimate density of other birds on the island. The extinct volcano is the northernmost island in the volcanic arc that extends between Sumatra and Myanmar (Bandopadhyay 2017). It is less than 700,000 years old and it became dormant in the Holocene (Bandopadhyay 2017). It is 130 km due east from north Andamans and 265 km from Myanmar, which is to the north and east of the Narcondam Island. The island rises abruptly to an elevation of about 710 m above sea level. Like typical stratovolcanoes, Narcondam is characterized by sharp, steep ridges and narrow ravines, making it extremely difficult to access various parts of the island. The littoral forests on the island are characterized by *Hibiscus tiliaceus*, *Pandanus odorifer*, *Ixora brunnescens* and *Guettarda speciosa*. The deciduous forests are characterized by *Bombax insigne*, *Gyrocarpus americanus* and *Ficus rumphii*. The evergreen forests are characterised by *Planchonella longipetiolata*, *Chionanthus* sp., *Syzygium clavatum*, *Caryota mitis*, and *Dysoxylum cyrtobotryum*. The understorey vegetation is dominated by *Mallotus resinosus* and *Actephila excelsa*.

**Methods: Tree watches**

The tree species included small-seeded figs (*Ficus benjamina*, *Ficus glaberrima, Ficus nervosa*, *Ficus rumphii*, and *Ficus geniculata*), small-seeded non-fig plants (*Aidia densiflora* and *Balakata baccata*), medium-seeded plants (*Anamirta cocculus*, *Chionanthus* sp. and *Codiocarpus andamanicus*) and large-seeded plants (*Caryota mitis* and *Endocomia macrocomia*) (Table S2). The tree watches generally started at 0600 h. In the case of fruiting trees observed in higher elevations, the fruit tree watches started between 0630-0650 h (n = 9 trees). Except for one individual each of *Aidia densiflora* and *Ficus nervosa*, all other trees were observed for around six hours (Table S2). The number of individuals observed per species varied between (1-16) (Table S2). Sixteen individuals of *Caryota mitis* were observed since the data was collected for another study focused on that species (Gopal et al. in prep.). The total effort invested in focal tree watches was 239 h 43 min. The effort was 143 h 28 min without including *Caryota mitis* (Table S2). Species with low detection probability may get underrepresented in the trail data. On the other hand, species with low detection probability were far less likely to be missed in tree watches. The chosen trees for tree watches provided good visibility of the entire canopy and the observer spent an extended duration just observing the focal fruiting tree. During the tree watch, we recorded species identity, the number of individuals, arrival and departure times of frugivores and fruit handling behavior to ascertain that the visitor was the seed disperser for the focal tree species following Naniwadekar et al. (2019*a*).

**Analysis: Frugivore densities**

For the Narcondam Hornbill, we had sufficient detections across the three elevation zones, therefore, we also estimated densities of hornbills across the different elevation zones to determine if the hornbill densities varied across the elevation. We truncated 5% of extreme distances (Thomas et al. 2010). To determine the model that best approximates the true detection function, we used the following combinations of key and adjustment terms: uniform key with cosine adjustments, half-normal key with cosine and Hermite polynomial adjustments and hazard-rate key with simple polynomial adjustments (Thomas et al. 2010). We used the Cramer-von-Mises test to assess the fit of the model with the data. Model with the least Akaike Information Criterion (AIC) value was selected.

**Modularity analysis**

We generated 100 null models using *null model 2* that implements the Monte Carlo procedure to determine the significance of the observed modularity (Bascompte et al. 2003). Stochastic nature of the algorithm may report different numbers of modules and modularity values in different runs. Therefore, we retained the configuration that had the maximum modularity value after running the modularity algorithm 25 times following Donatti et al. (2011).
