## Supplementary material for "Large frugivores matter more on an island: insights from island-mainland comparison of plant-frugivore communities": ESM Online Resource 2

**ELECTRONIC SUPPLEMENTARY MATERIAL 2**

**Table S1** Details of the 20 trails that were monitored to estimate bird abundance. We also report the number of visual detections of birds in each trail.

| **No.** | **Trail name** | **Elevation zone**  **(m ASL)** | **Trail length (km)** | **# of times trail walked** | **Total effort (km)** | **Asian Koel** | **Narcondam Hornbill** | **Imperial-Pigeons** |
| --- | --- | --- | --- | --- | --- | --- | --- | --- |
| 1 | Anchorage 0 | 0-200 | 0.9 | 1 | 0.9 | 0 | 3 | 2 |
| 2 | Basecamp to Pass | 0-200 | 0.56 | 5 | 2.8 | 2 | 14 | 6 |
| 3 | Coco Bay to Anchorage | 0-200 | 1.0 | 2 | 2.0 | 8 | 8 | 5 |
| 4 | Dead Narco Trail | 0-200 | 0.4 | 2 | 0.8 | 1 | 2 | 0 |
| 5 | Ekka Pahad | 0-200 | 0.4 | 1 | 0.4 | 2 | 1 | 0 |
| 6 | Pass to Coco Bay | 0-200 | 0.42 | 5 | 2.1 | 5 | 17 | 3 |
| 7 | Ravine Trail | 0-200 | 1.1 | 9 | 9.9 | 17 | 26 | 7 |
| 8 | Right Ridge 0 | 0-200 | 1.26 | 8 | 10.1 | 11 | 40 | 6 |
| 9 | Anchorage 200 | 200-400 | 0.8 | 2 | 1.6 | 3 | 6 | 1 |
| 10 | Loop Ridge | 200-400 | 0.7 | 3 | 2.1 | 2 | 9 | 2 |
| 11 | Mimusops 200 | 200-400 | 0.6 | 7 | 4.8 | 3 | 16 | 2 |
| **No.** | **Trail name** | **Strata**  **(m ASL)** | **Trail length (km)** | **# of times trail walked** | **Total effort (km)** | **Asian Koel** | **Narcondam Hornbill** | **Imperial-Pigeons** |
| 12 | Pukhri 400 | 200-400 | 0.75 | 2 | 1.5 | 1 | 5 | 3 |
| 13 | Right Ridge 200 | 200-400 | 0.96 | 5 | 4.8 | 8 | 31 | 7 |
| 14 | West of Pukhri Nala | 200-400 | 0.5 | 1 | 0.5 | 1 | 1 | 0 |
| 15 | Mimusops 400 | 400-700 | 0.44 | 5 | 2.2 | 1 | 5 | 0 |
| 16 | Mimusops 600 | 400-700 | 0.36 | 5 | 1.8 | 2 | 9 | 0 |
| 17 | Piper Trail | 400-700 | 0.9 | 1 | 0.9 | 0 | 7 | 0 |
| 18 | SE Top to Peak | 400-700 | 0.2 | 1 | 0.2 | 0 | 1 | 0 |
| 19 | South Ridge 600 | 400-700 | 0.6 | 1 | 0.6 | 1 | 1 | 0 |
| 20 | VM 400-600 | 400-700 | 0.8 | 2 | 1.6 | 3 | 5 | 1 |
|  | **Total** | **na** | **na** | **68** | **51.4** | **71** | **207** | **45** |

**Table S2.** Sampling details of tree watches that were conducted on the Narcondam Island

| **No.** | **Tree species** | **Number of individuals observed** | **Effort** |
| --- | --- | --- | --- |
| 1 | *Aidia densiflora* | 4 | 25 h 4 min |
| 2 | *Anamirta cocculus* | 1 | 6 h |
| 3 | *Balakata baccata* | 1 | 6 h |
| 4 | *Caryota mitis* | 16 | 96 h 15 min |
| 5 | *Chionanthus* sp. | 3 | 18 h |
| 6 | *Codiocarpus andamanicus* | 1 | 6 h |
| 7 | *Endocomia macrocomia* | 1 | 6 h |
| 8 | *Ficus benjamina* | 2 | 12h 10 min |
| 9 | *Ficus glaberrima* | 2 | 12 h |
| 10 | *Ficus nervosa* | 2 | 10 h 12 min |
| 11 | *Ficus rumphii* | 6 | 36 h 2 min |
| 12 | *Ficus geniculata* | 1 | 6 h |
|  | **Total** | **40** | **239 h 43 min** |

**Table S3.** (A) Summary of 27 plant species that were fruiting on Narcondam Island during our study period. Close relatives (in case target species was not found) that were used for the phylogenetic analysis. Fruit width of many species was measured in the field. For species which were not measured (*), fruit width information was obtained from the Flora of China (<http://www.efloras.org/flora_page.aspx?flora_id=2>). Plant species were classified as small-seeded if the seed size was < 5 mm, medium-seeded if it was 5-15 mm and large-seeded if it was > 15 mm in width. (B) Summary of plant species from the Pakke Tiger Reserve. Additional details can be found in Naniwadekar et al. (2019).

| **Species name** | **Species used in the phylogenetic analysis** | **Fruit width (mm)** | **Seed size** |
| --- | --- | --- | --- |
| *Aidia densiflora* | *Aidia racemosa* | 7.68 | Small |
| *Anamirta cocculus* | *Anamirta cocculus* | 13.11 | Medium |
| *Aphanamixis polystachya* | *Aphanamixis polystachya* | 24.83 | Medium |
| *Ardisia oxyphylla* | *Ardisia solanacea* | 9.7 | Medium |
| *Callicarpa arborea* | *Callicarpa arborea* | 3.73 | Small |
| *Caryota mitis* | *Caryota mitis* | 15.61 | Large |
| *Chionanthus* sp.* | *Chionanthus ligustrinus* | 14 | Medium |
| *Diospyros kurzii** | *Diospyros borneensis* | 20 | Small |
| *Discospermum abnorme* | *Discospermum polyspermum* | 15 | Small |
| *Ficus benjamina* | *Ficus benjamina* | 11.61 | Small |
| *Ficus glaberrima* | *Ficus drupacea* | 13.31 | Small |
| *Ficus microcarpa* | *Ficus microcarpa* | 6.5 | Small |
| *Ficus nervosa* | *Ficus nervosa* | 21.89 | Small |
| *Ficus rumphii* | *Ficus rumphii* | 11.06 | Small |
| *Ficus geniculata** | *Ficus superba* | 6 | Small |
| *Ficus virens** | *Ficus virens* | 8 | Small |
| *Ficus chartacea** | *Ficus chartacea* | 6 | Small |
| *Codiocarpus andamanicus* | *Gomphandra javanica* | 8.82 | Medium |
| *Endocomia macrocomia* | *Horsfieldia punctatifolia* | 28.67 | Large |
| *Ixora barbata** | *Ixora javanica* | 10 | Medium |
| *Lannea coromandelica* | *Lannea schweinfurthii* | 8.5 | Small |
| *Leea indica** | *Leea indica* | 7 | Small |
| *Mussaenda macrophylla** | *Mussaenda erythrophylla* | 12 | Small |
| *Planchonella longipetiolata* | *Planchonella obovata* | 27.5 | Large |
| *Pycnarrhena lucida* | *Pycnarrhena lucida* | 24.5 | Large |
| *Balakata baccata** | *Sapium glandulosum* | 12 | Small |
| *Schefflera* sp. | *Schefflera elliptica* | 2 | Small |

| **Species name** | **Species used in the phylogenetic analysis** | **Fruit width (mm)** | **Seed size** |
| --- | --- | --- | --- |
| *Aglaia spectabilis* | *Aglaia spectabilis* | 16.46 | Medium |
| *Antidesma montanum* | *Antidesma montanum* | 5 | Small |
| *Beilschmiedia* sp, 1 | *Beilschmiedia roxburghiana* | 14.76 | Medium |
| *Beilschmiedia* sp. 2 | *Beilschmiedia pendula* | 20.3 | Large |
| *Bischofia javanica* | *Bischofia javanica* | 9.92 | Medium |
| *Bridelia glauca* | *Bridelia glauca* | 8.82 | Small |
| *Callicarpa arborea* | *Callicarpa arborea* | 3 | Small |
| *Carallia brachiata* | *Carallia brachiata* | 9.38 | Medium |
| *Chisocheton cumingianus* | *Chisocheton cumingianus* | 23.34 | Large |
| *Cinnamomum bejolghota* | *Cinnamomum bejolghota* | 8.08 | Medium |
| *Dendrocnide sinuata* | *Dendrocnide sinuata* | 4.5 | Small |
| *Dysoxylum cauliflorum* | *Dysoxylum arborescens* | 12.67 | Medium |
| *Dysoxylum gotadhora* | *Dysoxylum parasiticum* | 21.36 | Large |
| *Dysoxylum procerum* | *Dysoxylum spectabile* | 19.64 | Large |
| *Ehretia wallichiana* | *Ehretia acuminata* | 7 | Small |
| *Elaeocarpus sphaericus* | *Elaeocarpus angustifolius* | 25.79 | Large |
| *Falconeria insignis* | *Sapium glandulosum* | 7.91 | Medium |
| *Ficus altissima* | *Ficus altissima* | 18.84 | Small |
| *Ficus benjamina* | *Ficus benjamina* | 16.26 | Small |
| *Ficus drupacea* | *Ficus drupacea* | 31.63 | Small |
| *Ficus geniculata* | *Ficus superba* | 6.94 | Small |
| *Ficus nervosa* | *Ficus nervosa* | 10.85 | Small |
| *Ficus obtusifolia* | *Ficus obtusifolia* | 12.53 | Small |
| *Ficus* sp. 1 | *Ficus erecta* | 12.12 | Small |
| *Heteropanax fragrans* | *Heteropanax fragrans* | 7.62 | Small |
| *Horsfieldia kingii* | *Horsfieldia punctatifolia* | 20.73 | Large |
| *Knema erratica* | *Knema laurina* | 15.16 | Medium |
| *Leea indica* | *Leea indica* | 7 | Small |
| *Litsea* sp. 2 | *Litsea panamanja* | 10.01 | Medium |
| *Livistona jenkinsiana* | *Livistona chinensis* | 27.75 | Large |
| *Micromelum integerrimum* | *Micromelum minutum* | 8.75 | Medium |
| *Olea dioica* | *Olea paniculata* | 11.1 | Medium |
| *Phoebe goalparensis* | *Phoebe angustifolia* | 12.32 | Medium |
| *Phoebe* sp. 1 | *Phoebe hungmoensis* | 10 | Medium |
| *Picrasma javanica* | *Picrasma javanica* | 10.62 | Medium |
| *Polyalthia simiarum* | *Polyalthia cerasoides* | 20.94 | Medium |
| *Prunus ceylanica* | *Prunus africana* | 21.92 | Medium |
| *Sloanea sterculiaceae* | *Sloanea sinensis* | 9.96 | Medium |
| *Sterculia villosa* | *Sterculia lanceolata* | 6.95 | Medium |
| *Syzygium* sp. 1 | *Syzygium cumini* | 8.73 | Medium |
| *Tetradium glabrifolium* | *Tetradium glabrifolium* | 4 | Small |
| *Vitex glabrata* | *Vitex glabrata* | 13 | Small |
| *Zanthoxylum rhetsa* | *Zanthoxylum acanthopodium* | 7.72 | Medium |

**Table S4.** Summary of frugivore species that were detected foraging on fruits on the Narcondam Island. Body mass and degree of frugivory (percent of fruits in the diet) is summarized below. Body mass and degree of frugivory information was obtained from Wilman et al. (2014).

| **Frugivore common name** | **Frugivore scientific name** | **Body mass (g)** | **Degree of frugivory** |
| --- | --- | --- | --- |
| Narcondam Hornbill | *Rhyticeros narcondami* | 697.17 | 100 |
| Pied Imperial-Pigeon | *Ducula bicolor* | 440.01 | 100 |
| Green Imperial-Pigeon | *Ducula aenea* | 545 | 90 |
| Asian Koel | *Eudynamys scolopaceus* | 194.92 | 80 |
| Common Hill Myna | *Gracula religiosa* | 192 | 60 |
| Daurian Starling | *Agropsar sturninus* | 49.4 | 20 |
| Eye-browed Thrush | *Turdus obscurus* | 62.6 | 60 |

**Table S5.** Estimates of detection probability, flock size, flock and individual densities and abundance and the associated coefficients of variation (CV), standard errors (SE) or 95% confidence intervals for the Narcondam Hornbill *Rhyticeros narcondami*, Asian Koel *Eudynamys scolopacea*, and Imperial-Pigeons *Ducula* spp. We estimated combined densities of the two Imperial-Pigeons (Green Imperial-Pigeon *Ducula aenea* and Pied Imperial-Pigeon *Ducula bicolor*) as we did not have sufficient (> 40) visual detections of each the two species.

| **Parameter** | **Species/group** | **Strata** | **Number of detections** | **Estimate** | **CV** | **SE** | **95% CI** |
| --- | --- | --- | --- | --- | --- | --- | --- |
| **Detection probability** | Narcondam Hornbill | Overall | 205 | 0.554 | 0.037 | 0.020 | - |
| **Detection probability** | Narcondam Hornbill | 0-200 m | 110 | 0.448 | 0.071 | 0.032 | - |
| **Detection probability** | Narcondam Hornbill | 200-400 m | 67 | 0.663 | 0.103 | 0.068 | - |
| **Detection probability** | Narcondam Hornbill | 400-700 m | 28 | 0.763 | 0.198 | 0.151 | - |
| **Detection probability** | Asian Koel | Overall | 70 | 0.674 | 0.101 | 0.068 | - |
| **Detection Probability** | Imperial-Pigeons | Overall | 45 | 0.292 | 0.138 | 0.040 | - |
| **Flock size** | Narcondam Hornbill | Overall | 205 | 1.745 | 0.068 | 0.118 | - |
| **Flock size** | Narcondam Hornbill | 0-200 m | 110 | 1.913 | 0.095 | 0.182 | - |
| **Flock size** | Narcondam Hornbill | 200-400 m | 67 | 1.578 | 0.039 | 0.062 | - |
| **Flock size** | Narcondam Hornbill | 400-700 m | 28 | 1.462 | 0.101 | 0.147 | - |
| **Flock size** | Asian Koel | Overall | 70 | 1.224 | 0.058 | 0.071 | - |
| **Flock size** | Imperial-Pigeons | Overall | 45 | 1.905 | 0.133 | 0.253 | - |
| **Flock density (per ha)** | Narcondam Hornbill | Overall | 205 | 0.862 | 0.115 | - | 0.681–1.092 |
| **Flock density (per ha)** | Narcondam Hornbill | 0-200 m | 110 | 0.845 | 0.166 | - | 0.586–1.220 |
| **Flock density (per ha)** | Narcondam Hornbill | 200-400 m | 67 | 1.062 | 0.207 | - | 0.666–1.692 |
| **Flock density (per ha)** | Narcondam Hornbill | 400-700 m | 28 | 0.692 | 0.294 | - | 0.374–1.278 |
| **Flock density (per ha)** | Asian Koel | Overall | 70 | 0.323 | 0.183 | - | 0.224–0.466 |
| **Flock density (per ha)** | Imperial-Pigeons | Overall | 45 | 0.198 | 0.212 | - | 0.130–0.302 |
| **Parameter** | **Species/group** | **Strata** | **Number of detections** | **Estimate** | **CV** | **SE** | **95% CI** |
| **Individual density (per ha)** | Narcondam Hornbill | Overall | 205 | 1.505 | 0.151 | - | 1.102–2.055 |
| **Individual density (per ha)** | Narcondam Hornbill | 0-200 m | 110 | 1.618 | 0.249 | - | 0.922–2.840 |
| **Individual density (per ha)** | Narcondam Hornbill | 200-400 m | 67 | 1.675 | 0.211 | - | 1.039–2.701 |
| **Individual density (per ha)** | Narcondam Hornbill | 400-700 m | 28 | 1.011 | 0.344 | - | 0.483–2.117 |
| **Individual density (per ha)** | Asian Koel | Overall | 70 | 0.395 | 0.203 | - | 0.263–0.595 |
| **Individual density (per ha)** | Imperial-Pigeons | Overall | 45 | 0.378 | 0.244 | - | 0.232–0.615 |
| **Abundance** | Narcondam Hornbill | Overall | 205 | 1026.3 | 0.151 | - | 751.4–1401.8 |
| **Abundance** | Asian Koel | Overall | 70 | 269.6 | 0.203 | - | 179.1–405.9 |
| **Abundance** | Imperial-Pigeons | Overall | 45 | 257.6 | 0.244 | - | 158.1–419.5 |

**Table S6.** Summary of degree, species strength, and weighted betweenness for the different plant and frugivore species. They have been arranged in descending order of degree. Species with highest values have been highlighted in bold.

| **Group** | **Species** | **Degree** | **Species Strength** | **Weighted Betweenness** |
| --- | --- | --- | --- | --- |
| Plant | *Ficus glaberrima* | 6 | 0.987 | 0.070 |
| **Plant** | ***Ficus rumphii*** | **6** | **2.613** | **0.930** |
| Plant | *Ficus benjamina* | 5 | 0.365 | 0 |
| Plant | *Ficus geniculata* | 5 | 0.606 | 0 |
| Plant | *Ficus microcarpa* | 4 | 0.237 | 0 |
| Plant | *Ficus nervosa* | 4 | 0.504 | 0 |
| Plant | *Ficus virens* | 4 | 0.587 | 0 |
| Plant | *Aidia densiflora* | 3 | 0.165 | 0 |
| Plant | *Balakata barbata* | 3 | 0.290 | 0 |
| Plant | *Anamirta cocculus* | 2 | 0.028 | 0 |
| Plant | *Ardisia oxyphylla* | 2 | 0.018 | 0 |
| Plant | *Callicarpa arborea* | 2 | 0.074 | 0 |
| Plant | *Chionanthus* sp. | 2 | 0.153 | 0 |
| Plant | *Mussaenda macrophylla* | 2 | 0.058 | 0 |
| Plant | *Schefflera* sp. | 2 | 0.037 | 0 |
| Plant | *Aphanamixis polystachya* | 1 | 0.011 | 0 |
| Plant | *Caryota mitis* | 1 | 0.094 | 0 |
| Plant | *Codiocarpus andamanicus* | 1 | 0.009 | 0 |
| Plant | *Discospermum abnorme* | 1 | 0.009 | 0 |
| Plant | *Diospyros kurzii* | 1 | 0.004 | 0 |
| Plant | *Endocomia macrocomia* | 1 | 0.024 | 0 |
| Plant | *Ficus chartacea* | 1 | 0.009 | 0 |
| Plant | *Ixora baccata* | 1 | 0.006 | 0 |
| Plant | *Lannea coromandelica* | 1 | 0.080 | 0 |
| Plant | *Leea indica* | 1 | 0.018 | 0 |
| Plant | *Planchonella longipetiolata* | 1 | 0.006 | 0 |
| Plant | *Pycnarrhaena lucida* | 1 | 0.008 | 0 |
| **Frugivore** | ***Rhyticeros narcondami*** | **22** | **17.003** | **1.00** |
| Frugivore | *Eudynamys scolopaceus* | 16 | 5.539 | 0 |
| Frugivore | *Gracula religiosa* | 8 | 1.717 | 0 |
| Frugivore | *Ducula aenea* | 7 | 1.360 | 0 |
| Frugivore | *Ducula bicolor* | 7 | 0.885 | 0 |
| Frugivore | *Turdus obscurus* | 3 | 0.492 | 0 |
| Frugivore | *Agropsar sturninus* | 1 | 0.004 | 0 |

**Table S7.** Coefficient table summarizing the output of the general linear model that examined the influence of plant type (*Ficus*, large-seeded plants, medium-seeded plants and small-seeded non-fig plants and abundance on the species strength (log_10_ transformed) of each of the 27 plant species. *Ficus* had significantly higher species strength values and the species strength was not influenced by plant abundance.

|  | **Estimate** | **SE** | ***t*** | ***P*** |
| --- | --- | --- | --- | --- |
| **(Intercept – Plant Type: Fig)** | -1.03 | 0.506 | -2.035 | 0.054 |
| **Abundance** | 0.013 | 0.015 | 0.875 | 0.391 |
| **Plant Type: Large-seeded** | -3.386 | 0.955 | -3.545 | 0.002 |
| **Plant Type: Medium-seeded** | -3.247 | 0.831 | -3.909 | <0.001 |
| **Plant Type: Small-seeded** | -2.308 | 0.707 | -3.266 | 0.004 |

**Table S8.** A The four modules identified using the Program MODULAR are shown below. Across the 25 runs, the composition of the modules remained constant. B The scientific names of frugivores and plant codes are provided. Each module has been given a unique color and same color codes have been provided for the scientific names.

| **Module 1** | | | **Module 2** | | | **Module 3** | | **Module 4** | | |
| --- | --- | --- | --- | --- | --- | --- | --- | --- | --- | --- |
| **Plant** | **Seed disperser** | | **Plant** | **Seed disperser** | | **Plant** | **Seed disperser** | **Plant** | **Seed disperser** | |
| ANCO | ASKO | | AIDE | PIPI | | APPO | NAHO | CHSP | CHMY | |
| AROX |  | | BABA | GIPI | | CAMI |  | FIGL | EBTH | |
| CAAR |  | | FIBE | DAST | | COAN |  | FISU |  | |
| FIAS |  | | FIMI |  | | DIAB |  | FIVI |  | |
| LEIN |  | | FINE |  | | DIKU |  | LACO |  | |
| SCSP |  | | FIRU |  | | ENMA |  | MUMA |  | |
|  |  | |  |  | | IXBA |  |  |  | |
|  |  | |  |  | | PLLO |  |  |  | |
|  |  | |  |  | | PYLU |  |  |  | |
| **Seed disperser** | | | | | **Plant** | | | | | |
| ASKO | | *Eudynamys scolopacea* | | | ANCO | *Anamirta cocculus* | | | COAN | *Codiocarpus andamanicus* |
| PIPI | | *Ducula bicolor* | | | AROX | *Ardisia oxyphylla* | | | DIAB | *Discospermum abnorme* |
| GIPI | | *Ducula aenea* | | | CAAR | *Callicarpa arborea* | | | DIKU | *Diospyros kurzii* |
| DAST | | *Agrospar sturninus* | | | FIAS | *Ficus asperrima* | | | ENMA | *Endocomia macrocomia* |
| NAHO | | *Rhyticeros narcondami* | | | LEIN | *Leea indica* | | | IXBA | *Ixora barbata* |
| CHMY | | *Gracula religiosa* | | | SCSP | *Schefflera* sp | | | PLLO | *Planchonella longipetiolata* |
| EBTH | | *Turdus obscurus* | | | AIDE | *Aidia densiflora* | | | PYLU | *Pycnarrhena lucida* |
|  | | | | | BABA | *Balakata baccata* | | | CHSP | *Chionanthus* sp |
|  |  |  |  |  | FIBE | *Ficus benjamina* | | | FIGL | *Ficus glaberrima* |
|  |  |  |  |  | FIMI | *Ficus microcarpa* | | | FISU | *Ficus superba* |
|  |  |  |  |  | FINE | *Ficus nervosa* | | | FIVI | *Ficus rumphii* |
|  |  |  |  |  | FIRU | *Ficus rumphii* | | | LACO | *Lannea coromandelica* |
|  |  |  |  |  | APPO | *Aphanamixis polystachya* | | | MUMA | *Mussaenda macrophylla* |
|  |  |  |  |  | CAMI | *Caryota mitis* | | | | |

**Figure S1.** Map showing the 20 trails monitored for estimating bird abundance. The 20 trails were spread across the elevation gradient. Eight trails in 0-200 m elevation strata and six each in 200-400 m and 400-700 m elevation strata. The yellow pin was our base camp and the blue pin is the Narcondam Peak. 1

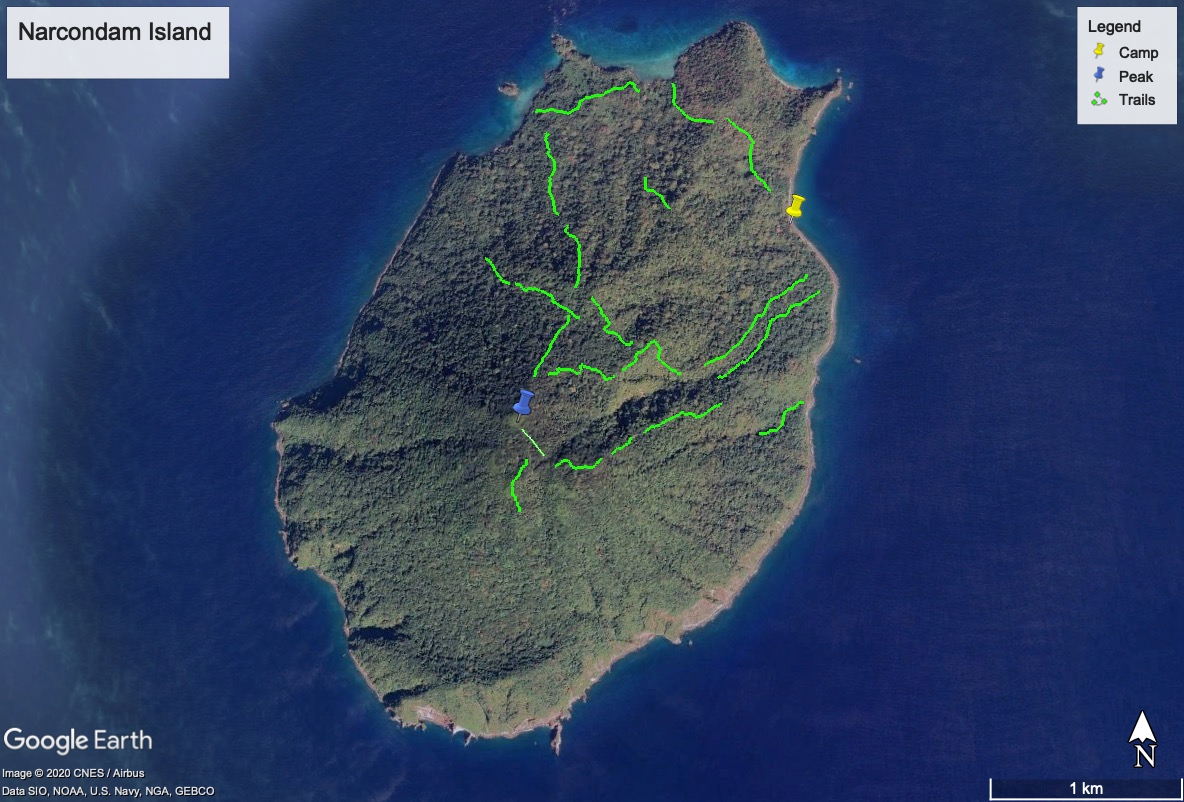

**Figure S2.** Interaction accumulation curve demonstrating that diversity of interactions had been adequately documented for the study period.

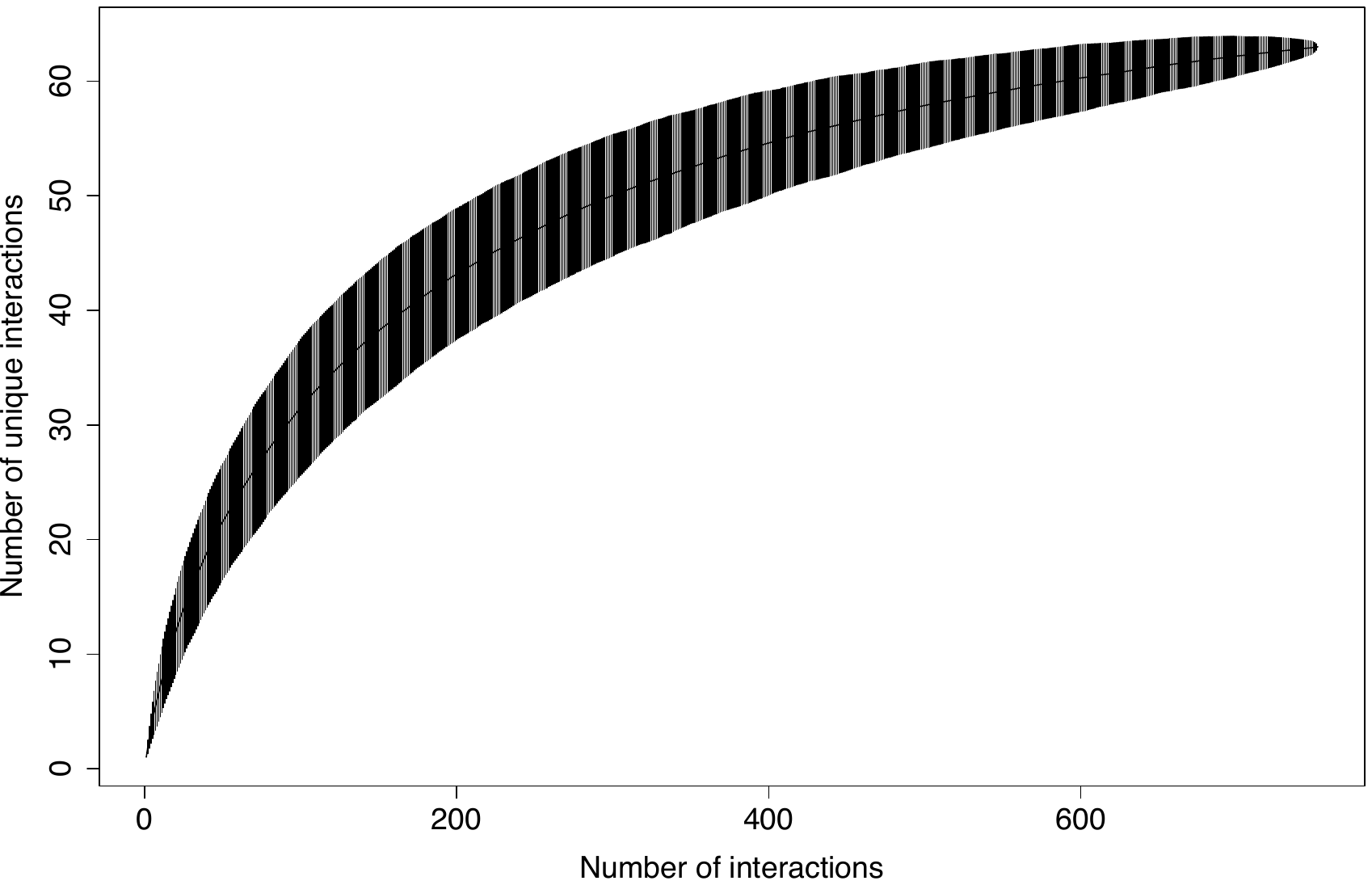

**Figure S3.** The network of plants and seed dispersers. Plants and frugivores are arranged in descending order of row and column totals. Frugivores and plants have been labelled following the L-R sequence of acronyms in the network. The Narcondam Hornbill (depicting frugivores) and *Ficus* (depicting plants) cartoons have been sketched by Sartaj Ghuman.

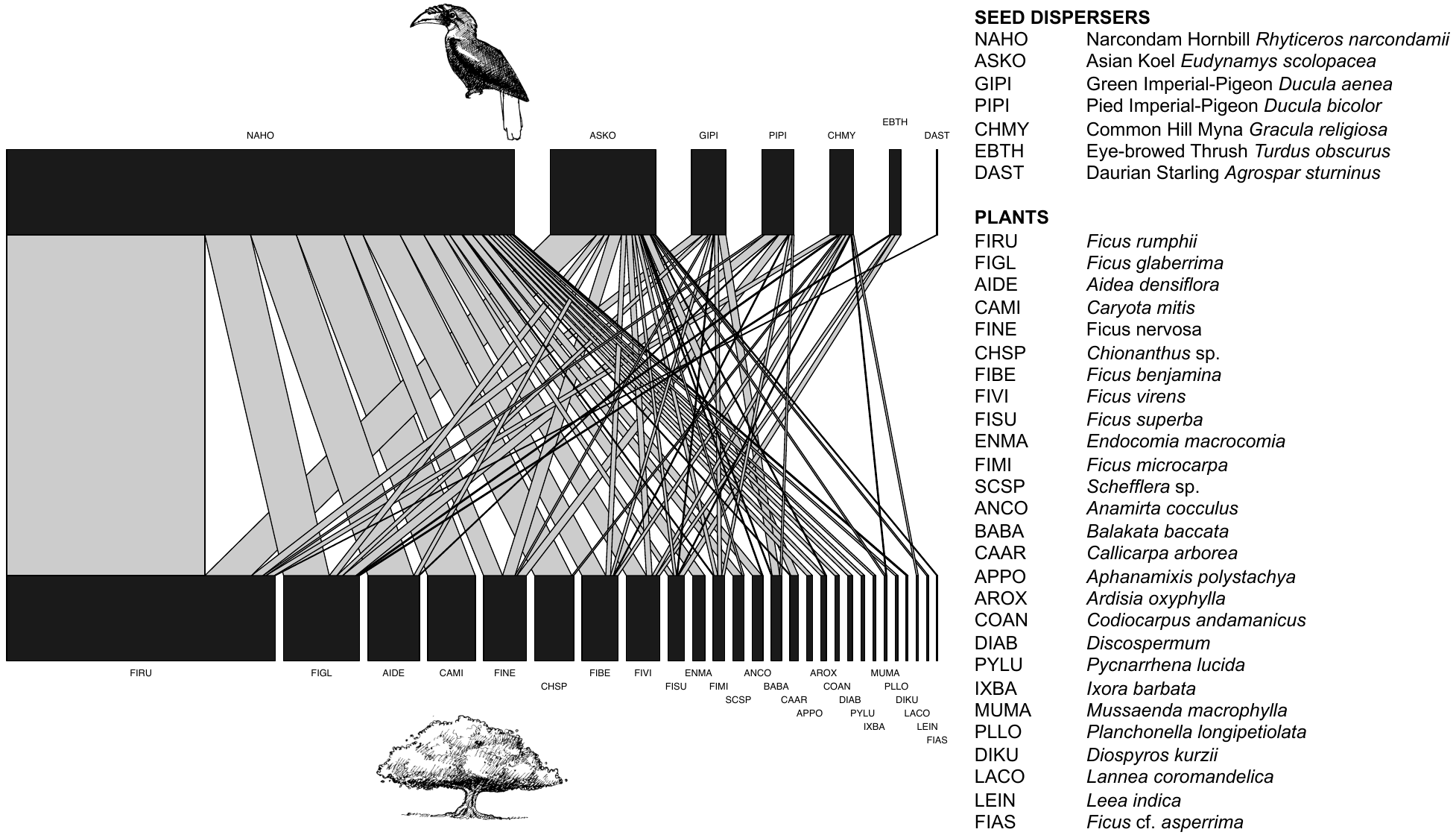
